# CRUX, a platform for visualising, exploring and analysing cancer genome cohort data

**DOI:** 10.1101/2023.07.25.550585

**Authors:** Sam El-Kamand, Julian M.W. Quinn, Heena Sareen, Therese M. Becker, Marie Wong-Erasmus, Mark J. Cowley

## Abstract

To better understand how tumours develop, identify prognostic biomarkers, and find new treatments, researchers have generated vast catalogues of cancer genome data. However, these datasets are complex so interpreting their important features requires specialized computational skills and analytical tools, which presents a significant technical challenge. To address this, we developed CRUX, a platform for exploring genomic data from cancer cohorts. CRUX enables researchers to perform common analyses including cohort comparisons, biomarker discovery, survival analysis, and create visualisations including oncoplots and lollipop charts. CRUX simplifies cancer genome analysis in several ways: (1) it has an easy-to-use graphical interface; (2) it enables users to create custom cohorts, as well as analyse precompiled public and private user-created datasets; (3) it allows analyses to be run locally to address data privacy concerns (though an online version is also available); and (4) it makes it easy to use additional specialized tools by exporting data in the correct formats. We showcase CRUX’s capabilities with case studies employing different types of cancer genome analysis, demonstrating how it can be used flexibly to generate valuable insights into cancer biology. CRUX is freely available at https://github.com/CCICB/CRUX and https://ccicb.shinyapps.io/crux (DOI: 10.5281/zenodo.8015714).

## INTRODUCTION

The complement of mutations present in every tumour is unique, with extensive molecular heterogeneity seen within and between tumour types. Among the myriad of mutations found in a tumour, relatively few actively drive its genesis and development. This excess of passenger mutations makes it challenging to determine the importance of individual mutations in cancer genome datasets. Identifying, cataloguing and inferring the functional impact of somatic single nucleotide variants (SNVs), short insertions and deletions (indels) and copy number variants (CNVs) has thus been a major research focus for the last decade. Next-generation sequencing (NGS) methods make possible the study of such mutations across many individuals in both a retrospective research context and now, increasingly, in a clinical context. This large and growing body of available cancer mutation data has obvious value to researchers seeking to understand the diagnostic, prognostic, and therapeutic consequences of individual variants. Steep falls in sequencing costs will also make possible greater use of NGS in clinical practice, however, analysing mutations in such datasets still requires strong technical skills and a working knowledge of many bioinformatics techniques. Lowering these technical barriers would make study of NGS datasets far easier, facilitate discovery of novel cancer gene drivers, and encourage both hypothesis- and curiosity-driven exploration of datasets.

Several online tools for exploring cancer NGS cohorts using graphical interfaces are already available, including the St. Jude Cloud (1), cBioPortal (2), GenePattern (3), Mutalisk (4), and the cancer virtual cohort discovery analysis platform (CVCDAP) (5) (Supplementary Table S1). However, important challenges remain unsolved. Many of these platforms are primarily designed for exploration of large public datasets generated by initiatives like The Cancer Genome Atlas (TCGA) (6) and Pan-Cancer Analysis of Whole Genomes (PCAWG) (7). It often remains difficult to leverage the full capabilities of tools like cBioPortal when studying unpublished, user-generated datasets, as this requires the technical knowledge to set up a local instance of the tool, or a willingness to add datasets to centralised repositories prior to exploration. Even tools which can be used out of the box to study user datasets (e.g. CVCDAP and GenePattern) still require data to be uploaded to an external server, with no viable alternatives available (for those without programming skills) that would mitigate data privacy concerns. Another key limitation of the current ecosystem is insufficient inter-operability. The most comprehensive graphical tools available for exploring mutational signatures, identifying driver mutations and annotating variants are all completely independent platforms. To use these tools, researchers must first convert their datasets into appropriate file formats. After analysis, processing is also frequently required to extract cohort-level insights. This pre- and post-processing must be repeated for each specialised analysis run and costs researchers time or may result in underuse of valuable tools due to the burden of entry. Due to these many issues, the current landscape of cancer cohort analysis tools would benefit from a new addition which can 1) run identical analyses on both private and public datasets, 2) import and export data in formats compatible with independently maintained software tools, 3) easily be installed locally to mitigate data privacy concerns and 4) offer an intuitive graphical interface.

We therefore developed CRUX as an intuitive analysis platform to address these issues after experiencing first-hand the difficulties of researchers and clinicians trying to extract actionable information from NGS cancer datasets. CRUX provides a rich genomic data exploration environment that integrates tertiary analysis and visualisation tools. Many of these tools can be used offline, and cancer genome datasets can be loaded, manipulated, and studied using a dashboard-style user interface. Data analysis and visualisation in CRUX employs many of the tools provided by *maftools*, a popular cancer cohort data exploration R package (8). *Maftools* supports a range of cancer cohort data manipulations and visualisations, such as oncoplot creation, cohort comparisons and identification of mutated pathways. However, analysis packages such as *maftools* still require programming skills lacking in many researchers. CRUX circumvents these issues to provide a simple and secure user application. It also provides direct access to genomic and clinical data from 11,043 tumour samples from 57 pre-compiled TCGA and PCAWG (8, 9) cohorts (8-10). Furthermore, CRUX enables creation of virtual cohorts by merging and sub-setting datasets based on clinical or molecular features, with a ‘multiple cohort’ mode that allows comparative studies of cohorts or sub-cohorts. There are also gold-standard and cutting-edge web-based analysis platforms a researcher may need to employ to search for a signal of interest (11). CRUX facilitates use of these external tools (see Results section), which improves the breadth of available analysis that can be performed by giving access to new (and constantly emerging) state-of-the-art tools.

In summary, CRUX has been designed to encourage efficient and dynamic dataset exploration that can follow a developing line of enquiry to obtain novel (and serendipitous) research findings in cancer mutations. Here, we have used CRUX in a series of small studies, not primarily intended to generate novel clinical insights but rather to demonstrate the capabilities of CRUX in common analysis scenarios.

## MATERIALS AND METHODS

### Data Analysis

All analyses presented here were either produced using CRUX or through CRUX-accessed platforms as described in the sections below and the instructions available in the ‘external tools’ module.

### Tool Development

CRUX was written in R version 4.0.3. The graphical interface was built using Shiny, shinyWidgets, and shinydashboard. The core workspace provides a graphical interface for analyses and visualisations from the *maftools* package version 2.12.0 (8). Additional visualisations were added using ggplot2 and ggVennDiagram libraries. All tables are rendered using the DT library. All code is packaged in a golem framework, allowing the CRUX shiny app to be distributed as an R package.

### Software availability

CRUX is freely available as a shiny app at https://ccicb.shinyapps.io/crux/ or can be downloaded from GitHub at https://github.com/CCICB/CRUX under a GNU General Public License v3.0 open-source license, DOI: 10.5281/zenodo.8015714, https://opensource.org/licenses/GPL-3.0. CRUX can be installed as an R package on any R-compatible operating system. CRUX functionality has been tested on Windows 11 and macOS. A manual is available at https://crux-docs.readthedocs.io/en/latest/.

## RESULTS

Cancer genome cohort datasets are generated from tumour samples that have undergone NGS and subsequent processing with dedicated mutation detection tools, usually in standardised analysis pipelines. Typical investigations of such cohort genomic data might include the following steps: 1) identify somatic cancer mutations; 2) identify driver mutations enriched in the cohort; 3) identify molecular subtypes of disease; 4) identify mutational signatures, 5) identify associations between molecular and clinical features, e.g., patient survival; and 6) identify biomarkers that predict therapeutic responses. CRUX is an integrated platform for performing such investigative analyses on genomic datasets (Fig. 1).

**Figure 1.**
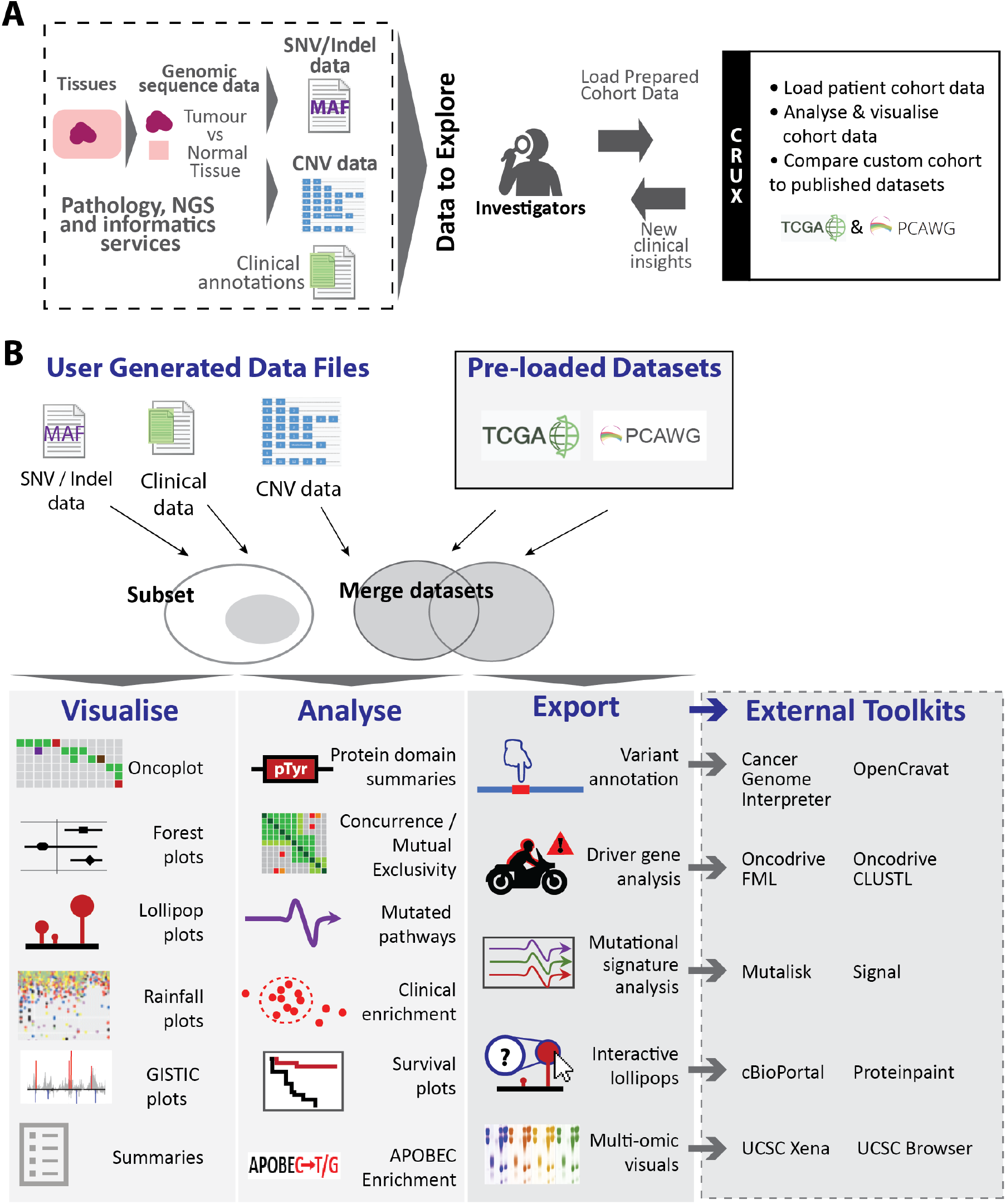
Tumour NGS data and CRUX. **A)** Patient tissue or cell samples undergo whole genome sequencing and informatics processing to prepare files describing tumour SNVs and indels (MAF format), copy number variations (CNV; GISTIC2 format), and clinical metadata. These can be loaded in CRUX to study patient tumour mutations, disease subtypes, signatures, and potential biomarkers of drug responses. **B)** CRUX workflows and modules. Imported data or cancer genome datasets from TCGA and PCAWG pre-loaded into CRUX can be used. Pre-loaded and user-imported data (subsetted or merged as required) are explored using *maftools*-based analyses with modules indicated. Export functions also facilitate use of many other toolkits (indicated) accessed through a new window or browser tab. Supplementary Table S2 describes available modules and toolkits.

### Design and use of CRUX for cancer genome cohort exploration

Somatic mutations are identified through comparison of patient NGS data from tumour and normal tissues. Common file formats for somatic SNV and indel data are mutation annotation format (MAF) and variant call format (VCF); CRUX directly supports importing MAF files for custom cancer NGS datasets, and provides instructions for converting VCFs to MAF format (12). CRUX provides a convenient graphical user interface (GUI) for data exploration, visualisation, and analysis by powerful computational modules. CRUX modules, functionalities and use are illustrated in Fig. 1 and Supplementary Fig. 1, with further details in Supplementary Table S2.

### Use of external toolkits for data analysis

CRUX provides access to external toolkits and platforms, i.e., resources that are available online (Fig. 1B) including tools that are specialised for variant annotation, mutational signature analysis and driver gene identification; many of these are accepted as benchmark analytical tools. CRUX reformats the datasets, exports them in the required format for the external tool, then opens the platform site where formatted files can be imported, and outputs and charts generated. Examples are given in Short Study 1, which utilises both internal modules and oncodriveFML (13). Similarly, Short Studies 3 and 4 use cBioPortal, Cancer Genome Interpreter, Mutalisk and Signal online resources (4, 11, 14, 15). These studies illustrate how use of internal and external resources can be interwoven to maximise utility.

### Custom cohort dataset creation

Merging and splitting of datasets to create virtual cohorts is useful for comparing molecular and clinical characteristics of different groups of patients. Such exploration is supported by CRUX, allowing sub-groups to be defined based on either patient-level clinical data or mutational profile. As an example, in Supplementary Fig. 2 TCGA subsets of breast cancers were created based on well-established breast cancer features, estrogen (ER) and progesterone (PR) using the data sub-setting and merging utilities.

**Figure 2.**
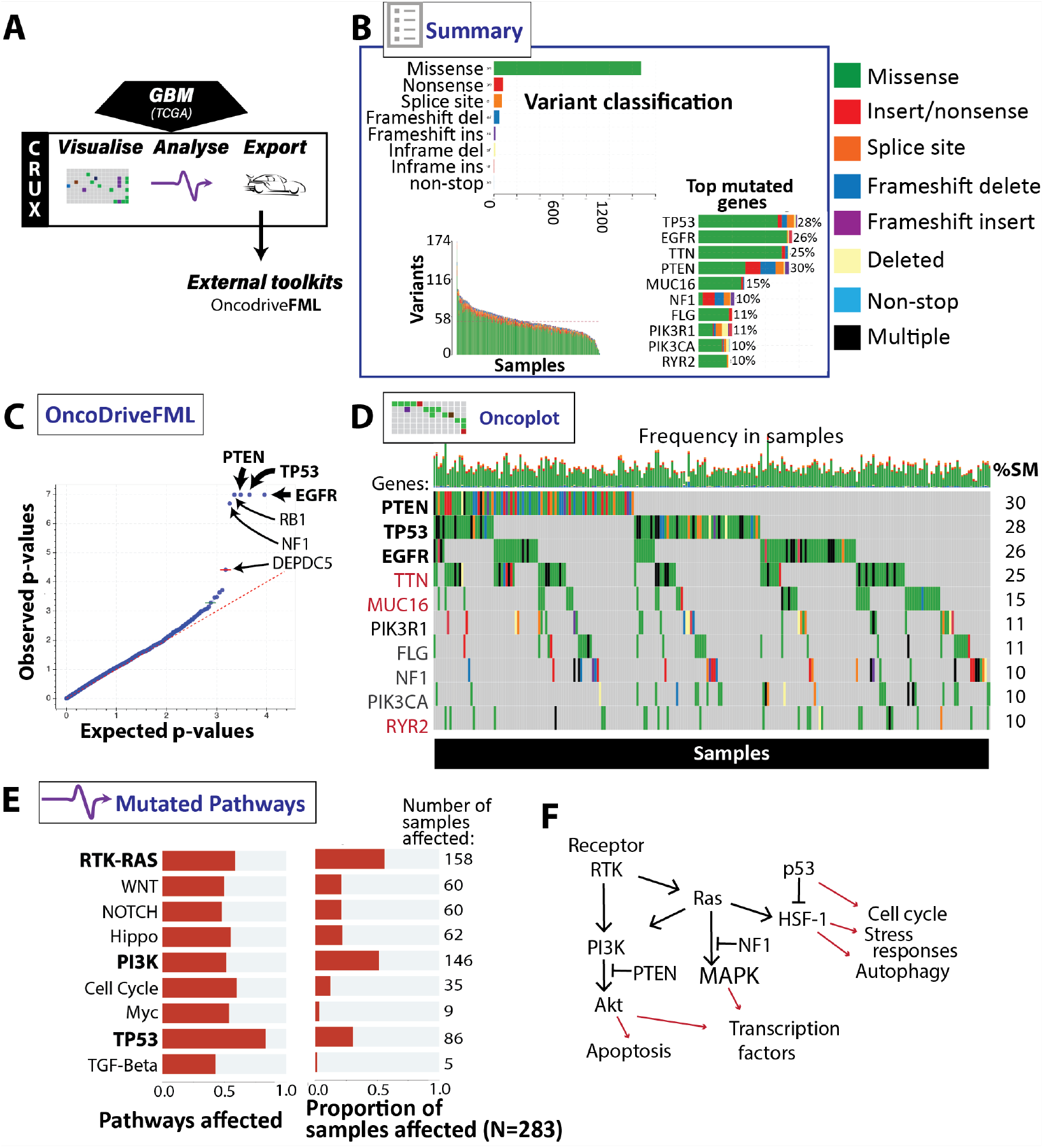
Identifying putative oncogenes in a Glioblastoma (GBM) cohort using CRUX. (A) CRUX modules used for TCGA GBM datasets. (B) Summary module enabled review of cohort mutation profiles with graphical outputs: prevalence of mutation types (top), number of mutations (bottom left) and mutation frequency by gene (bottom right). Mutation type key on right. (C) External tool OncoDriveFML identified genes selected in cancer (bold), including known GBM driver genes *EGFR, TP53* and *PTEN*; (D) Oncoplot module showed genes mutated in cancer samples, and frequency and type of mutations across samples; mutations clustered in *EGFR, TP53*, and *PTEN*. ‘%SM’= % samples with indicated gene mutated. Dubious candidates filtered out by CRUX shown in red. Note gene ordering is by number of samples with the gene mutated, not total mutations, as in B. (E) Visualisation of frequently mutated pathways in GBM, notably TP53, PI3K and RTK-RAS pathways. (F) Typical signal interactions of key gene products noted above. Ligand binding of receptor such as EGFR activates receptor tyrosine kinase (RTK) and thence the PI3K/Akt pathway (inhibited by PTEN) and Ras, which activates PI3K (and so NF1 inhibited MAPK pathways) and heat shock factor-1 (HSF-1), indirectly inhibited by p53 (TP53 product). These affect cell functions such as those indicated.

### CRUX short studies

Here, to demonstrate the utility of CRUX we describe six short studies below. Five used the CRUX single cohort analysis tools and one employed the ‘cohort comparison’ mode that investigates differences between cohorts or cohort sub-groups. These studies demonstrate that established features of cancers can easily be identified in the datasets, so researchers can either make similar discoveries in their own data or use published data in novel ways in their work. These studies returned outcomes or conclusions comparable to outcomes reported in published work, as indicated.

### Short study 1 (single cohort mode): Identify putative driver oncogenes from glioblastoma multiforme (GBM) tumours

#### Rationale

A major objective in cancer genome exploration is to identify cancer driver genes that, when mutated, promote some aspect of cancer initiation or progression. As most tumours accumulate thousands of mutations it is challenging to distinguish drivers from the many passenger mutations. While early efforts simply identified genes with most somatic mutations, algorithms have matured in the last 10 years and now correct for long, highly expressed, late-replicating genes (16) and filter out genes under minimal selective pressure, which corrects the previous artificially inflated driver gene estimates, as reviewed by Martinez-Jimenez et al (17). Here, to identify candidate GBM gene driver mutations we visualised, explored, and filtered genes bearing mutations from TCGA (6, 18) using CRUX.

#### Dataset

The TCGA glioblastoma multiforme (GBM) dataset (n = 283) pre-packaged with CRUX (see Materials and Methods).

#### Analysis and observations

Selection of the pre-packaged TCGA dataset in the ‘Cohort Summary’ mode of CRUX launched several mutation visualisations, including a summary of cohort mutation profiles. Modules used in this study (Fig. 2A) included Summary and Oncoplot visualisation modules, followed by analyses using Mutated Pathways and the external OncoDriveFML tool (13).

The Summary module detected a high frequency of mutations (mainly missense) in *PTEN, TP53* and *EGFR*, genes already identified as commonly mutated in the TCGA dataset (Fig. 2B), as well as *TTN*. To identify driver genes in GBM, we used CRUX to re-format and upload the data to OncoDriveFML, which again identified *PTEN, TP53, EGFR* as key driver genes (Fig. 2C). *NF1*, ranked 8^th^ by mutation frequency alone, has a similar p-value to the other driver genes. Note the titin gene (*TTN*) is large, allowing it to accumulate many passenger mutations, which OncoDriveFML correctly recognised as unlikely to be genuine driver mutations. Since other such false positive genes repeatedly appear in cancer genome analyses, due to large length or high somatic mutation rates by mechanisms unrelated to cancer, CRUX provides a simple filter option (‘Filter Dubious Genes’) to exclude these genes from further analysis. Exclusions are based on gene filters compiled from literature (16, 19) into the somaticflags R package. This CRUX option removes *TTN, MUC16 and RYR2* from all subsequent results.

Oncoplots are a popular visualisation for identifying patterns of gene co-mutation, here revealing it is rare for patients to have driver mutations in all three of *TP53, PTEN* and *EGFR*, and that different molecular subtypes exist (Fig. 2D). As most genes operate in pathways, we examined this with the Mutated Pathways module. This identified several affected pathways (Fig. 2E, F), notably TP53, PI3K and RTK/RAS pathways, all now with established influences in GBM. Generating oncoplots for each of these three pathways (Supplementary Fig. 3) reveals contributions of additional genes. For example, the RTK-RAS pathway is driven by *EGFR, NF1* with additional mutations in *PDGFRA*, in a mutually exclusive pattern (20). Analysis of the *PI3K* pathway alterations reveals *PTEN* as predominant driver (20), with additional contributions from *PIK3R1* and *PIK3CA* mutations, which tend to be mutually exclusive (Supplementary Fig. 3).

**Figure 3.**
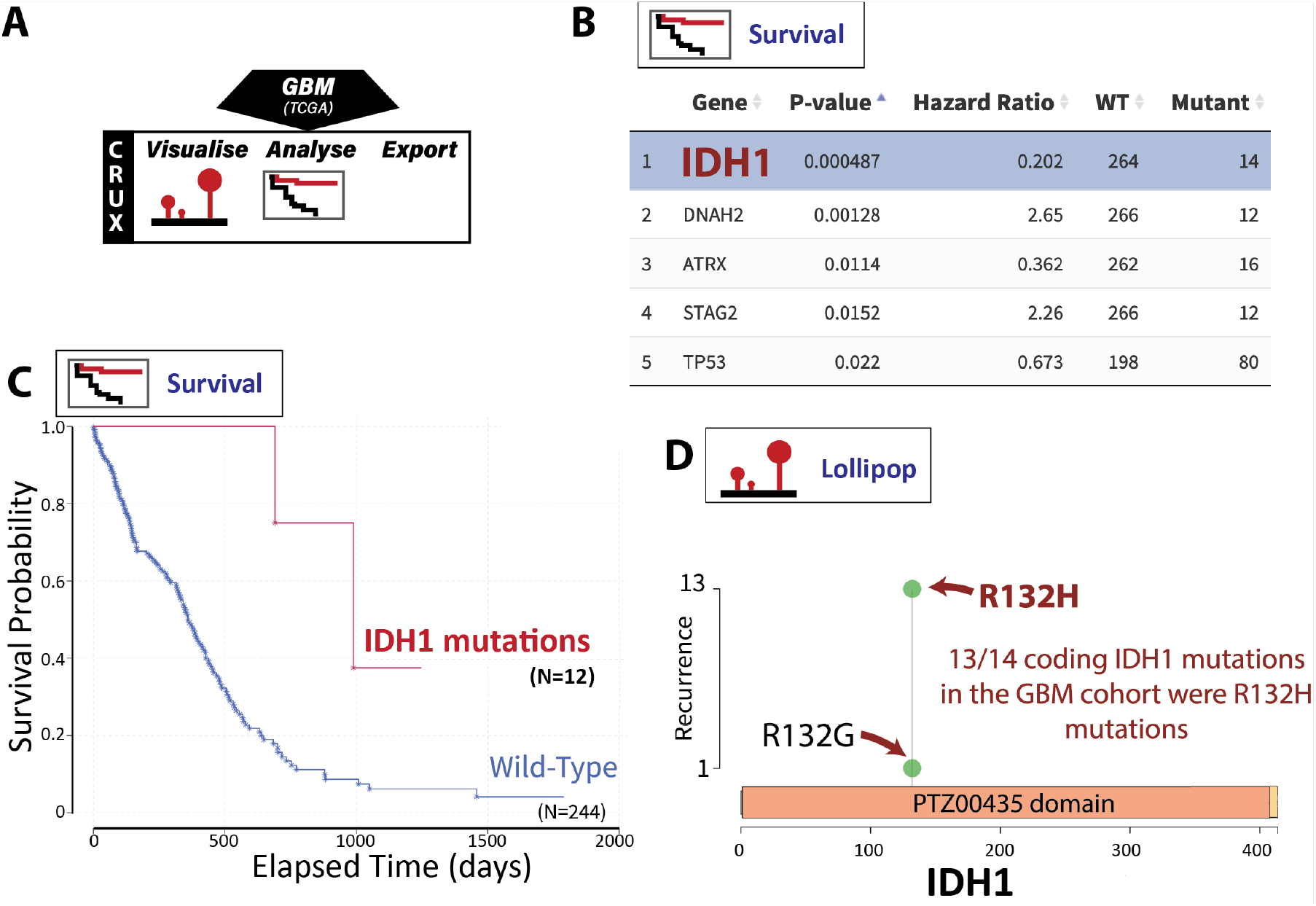
Finding prognostic biomarkers associated with longer patient survival in GBM. (A) The lollipop and survival plot modules were utilised in CRUX. (B) The Survival module identified several genes associated with better (hazard ratio <1) or worse survival (hazard ratio >1), with *IDH1* mutations being strongly associated with improved survival. (C) Kaplan Meier plot comparing the survival of patients with or without *IDH1* mutations. (D) Lollipop plots show *IDH1* mutations are focused at R132H/G, now a well-established prognostic biomarker.

*Outcome:* GBM driver mutations were identified by well-established driver selection software and interactions between these genes noted by Pathway Enrichment analysis and oncoplot visualisations. These findings are concordant with prior analyses of TCGA GBM cohort (21), but here were identified using no programming or command line tools.

### Short Study 2 (single cohort mode): Identify biomarkers associated with patient survival by integrating genome molecular alterations with clinical data

#### Rationale

Another major goal of cancer genomics is to find biomarkers associated with different prognoses and therapeutic responses. Here, we employed patient survival data to identify mutations that associate with survival in GBM patients.

#### Dataset

The inbuilt GBM cohort dataset (n = 283) from TCGA, as in short study 1.

#### Analysis and observations

The TCGA GBM cohort was selected in CRUX and the mutation data, with matched clinical annotations, was automatically loaded (Fig. 3A). To identify mutation biomarkers associated with survival time, CRUX’s Survival Analysis module was selected. For each mutated gene, CRUX performs a two-group survival analysis, calculating hazard ratios using a Cox proportional hazard model and p values by log-rank tests of Kaplan-Meier survival curves. The results are presented as a table showing the strongest potential prognostic markers (Fig. 3B). By these criteria the mutation biomarker associated most strongly with increased survival was seen in *IDH1* (p-value 0.001); Kaplan-Meier analysis confirming GBM tumours with mutated *IDH1* (isocitrate dehydrogenase isozyme) have better prognosis with mortality hazard ratio of 0.147 at 1000 days (p=0.0018, by log-rank tests of Kaplan Meier survival) (Fig. 3C). Further investigation of the location of the *IDH1* mutations within the gene using the lollipop module revealed that all mutations occurred at the R132 codon, indicating this is a hotspot mutation (Fig. 3D), indeed it is an established GBM prognostic biomarker and drug target.

#### Outcome

In this GBM dataset strongest associations with patient survival were seen with *IDH1* R132 mutations compared to patients with tumours bearing wild type *IDH1. IDH1* R132 mutation is previously described as a prognostic marker in GBM (22). This may be linked to reduced NAPH-related metabolism that sensitises cells to chemotherapy (23), as well as altered gene methylation patterns and oncogene activity (23, 24). This case study demonstrates CRUX functionality and utility in identifying such markers.

### Short study 3 (single cohort mode): Identification of candidate driver mutations linked to therapeutic responses in thyroid cancer

#### Rationale

Visualising location and clustering of mutations within a gene can help identify potential drivers of tumour progression; online resources can also help identify whether a particular mutation affects sensitivity to a targeted therapy.

#### Dataset

The Thyroid Cancer (THCA) dataset, consisting of 496 samples with whole genome sequencing data.

#### Analysis and observations

CRUX modules and tools used for this THCA dataset are shown in Figure 4A. Using oncoplots and the OncoDriveCLUSTL external driver gene analysis toolkit indicated *BRAF*, the Ras genes (*HRAS, NRAS, KRAS*) and the splicing regulator *RBM10* as key drivers in THCA (data not shown). To visualise the mutations in *BRAF* in THCA, the export module was used to reformat data for compatibility with the cBioPortal mutation mapper to produce interactive lollipop visualisations that revealed *BRAF* V600E mutations as common in these patient samples (Fig. 4B). Annotation tracks provided by the platform indicate that this variant is 1) at a known cancer-hotspot from the Cancer Hotspots and OncoKB resources (25); 2) in the middle of a tyrosine kinase protein domain; and 3) flanked by protein phosphorylation sites. Similarly, the mutations in *HRAS, NRAS, KRAS* and *RMB10* were all at well-established mutation hotspots (data not shown).

**Figure 4.**
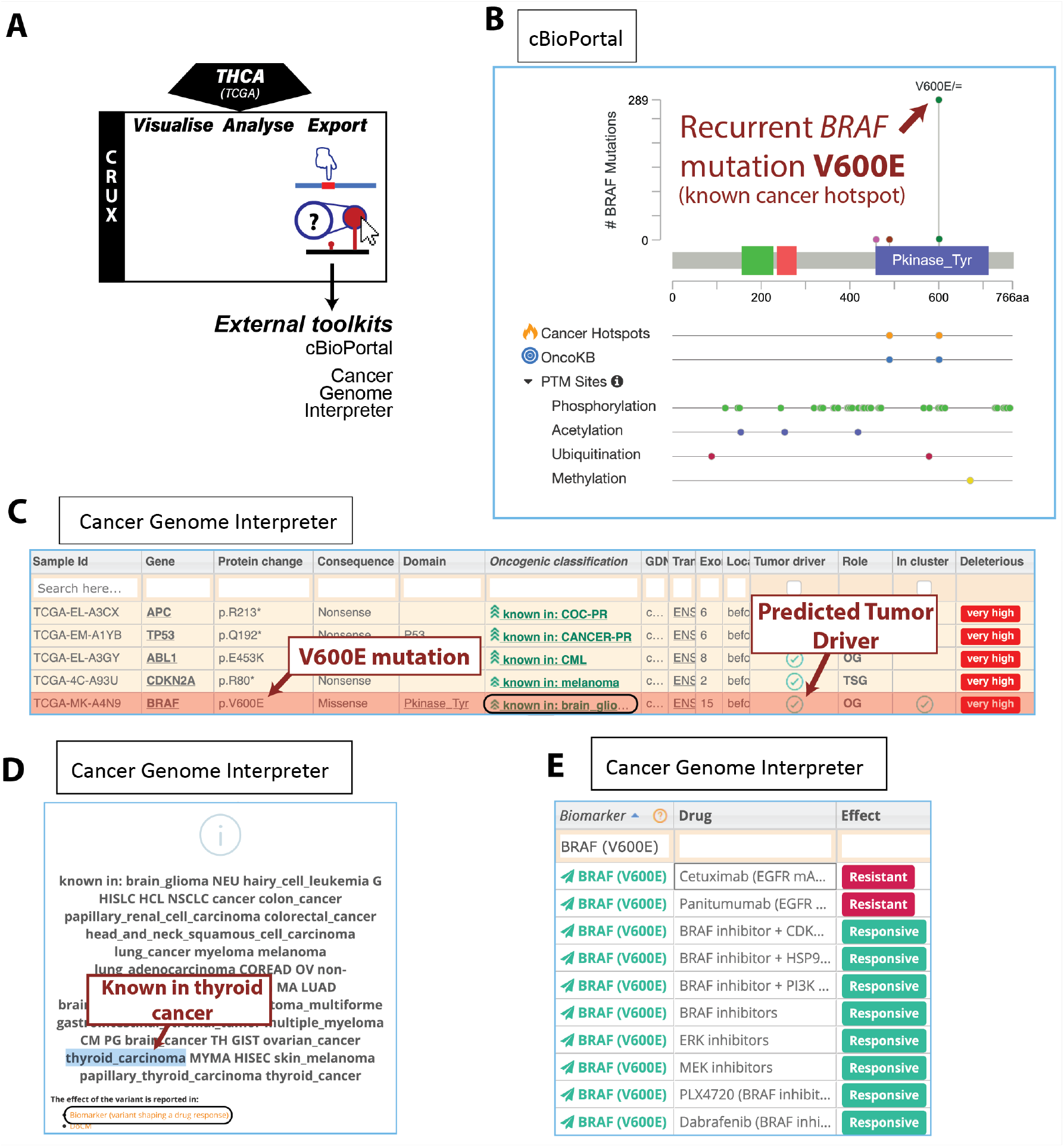
Characterising *BRAF* mutations as thyroid cancer drivers. (A) Modules used for short study 3: the study used TCGA thyroid carcinoma (THCA) cohort data pre-packaged in CRUX, with data exported to cBioPortal MutationMapper and Cancer Genome Interpreter (CGI) to facilitate variant-level interpretation. (B) Interactive lollipops are provided by cBioPortal MutationMapper, which provides annotations from various resources, including Cancer Hotspots, OncoKB and post-translational modifications. (C, D) CGI annotations indicated that *BRAF* V600E is an activating mutation known in THCA and several other cancers. (E) Sensitivity and resistance of *BRAF* V600E mutation carriers to clinical and pre-clinical therapeutics.

These observations motivated further investigation to determine whether *BRAF* V600E is commonly seen in thyroid cancer. The export module was once again used to produce formats permitting variant annotation by the Cancer Genome Interpreter (CGI), an online platform to guide variant-level interpretation of oncogenicity and therapeutic actionability (15). CGI indicated that 1) *BRAF* is a known oncogene (Figure 4C); 2) V600E is a common mutation seen in many types of cancers, including thyroid carcinoma (Fig. 4D); and 3) there are several approved targeted therapies for BRAF-mutant thyroid cancer (Figure 4E). *TTN* was shown to be mutated at high frequency in THCA but (similar to short study 1) were classified by CGI as a ‘predicted passenger’ (Supplementary Fig. 4). The *BRAF* V600E mutation is enriched in the conventional papillary thyroid cancer subtype, driving a more aggressive phenotype with higher risk of recurrence than patients without the mutation.

#### Outcomes

*BRAF* mutations were common in this THCA cohort and associated with features that suggest that such BRAF mutations are likely gene drivers and therapeutic targets. Despite several multi-tyrosine kinase and BRAF-specific inhibitors being FDA and EMA approved for BRAF-mutant THCA tumours they are less sensitive to these drugs than melanoma, rapidly developing resistance and new treatment strategies continue to be evaluated, including combination therapies (26).

### Short Study 4 (single cohort mode): Mutation signature analysis of cohort data

#### Rationale

Mutation signature analysis can provide mechanistic insights to the tumour development, indicating exposure to environmental stimuli (e.g., ultraviolet radiation or smoking) and the genome-wide impact of inherited and acquired driver mutations (e.g., damage to DNA repair pathways). Rich catalogues of mutation signatures have been developed but since these are constantly evolving (as is understanding of the tumour aetiology to which those signatures are linked), we would recommend the use of online analysis platforms to study them, which CRUX facilitates.

#### Dataset

We created a new dataset in CRUX by importing published variant calls from a previous study of 30 lung tumours sequenced with deep multi-region whole genome sequencing (WGS), merging this with the associated clinical data (27). The patients in this cohort were a range of current, former, and non-smokers, and the tumour biopsies were from paired primary and metastatic tumour biopsies.

#### Analysis and Observations

We used the CRUX mutational signature export module to facilitate analyses by two web-based analysis platforms, Mutalisk (4) and Signal (11) (Fig. 5A). CRUX formatted and exported the data in the correct format for these platforms, then we determined the mutational signatures by screening against all PCAWG (V3) signatures. Results from the Mutalisk analysis were imported back into the CRUX ‘mutational signature module’ for cohort-level visualisation and integration with clinical annotations.

**Figure 5.**
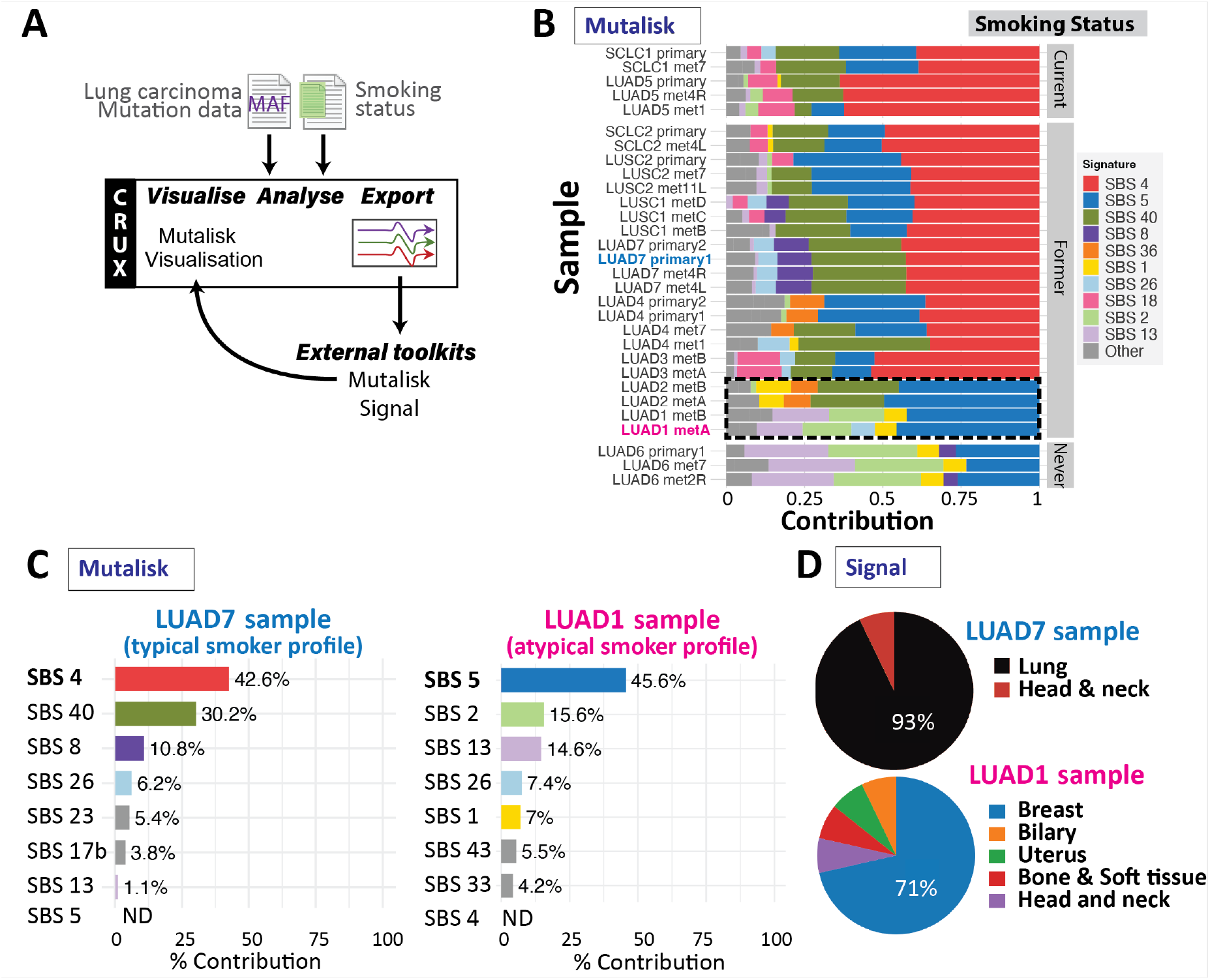
Identifying smoking-related mutation signatures, using a published lung cancer dataset from Leong et al. (27). (A) Workflow through CRUX using the Mutational Signature module that formatted and exported data to Mutalisk (4) and Signal (11); for each patient sample these external platforms were used to identify and quantify single bases substitution (SBS) mutational signatures from the COSMICv3 signature catalogue. (B) A cohort visualisation generated by importing Mutalisk results into the mutational signature module of CRUX shows their SBS signatures (indicated by colour) and their contribution to the sample mutation profile. Data from individual samples are shown (one per composite line) clustered into current smokers (top group), former smokers (middle group) and never smokers (bottom group); samples are included from both primary and metastases (‘met’) for most patients. Most current and former smokers (but not never-smoked controls) exhibit the SBS4 (dark blue) smoking associated-signature. However, tumours of some former smokers lack this SBS4 signature (black dashed box). (C) Mutalisk signature analysis output confirmed that sample LUAD1_metA derived from a former smoker lacked the smoking-associated SBS4 signature (‘ND’), typical of tumours in smokers such as LUAD7; this aligns with observations of Leong et al. (27). (D) Comparison of mutational signatures to samples in the Signal database reveals tumours lacking SBS4 (including LUAD1) are atypical, with low cosine similarity to confirmed lung carcinomas, while the LUAD7 profile (typical smoker) is seen in lung and head and neck cancers in the Signal database.

As expected from the original study (27), Mutalisk analysis indicated that primary and metastatic lung tumour tissue from most current and former smokers had a prominent SBS4 signature (Fig. 5B), associated with exposure to carcinogens in cigarette smoke. However, tumours from two former smokers (LUAD1 and LUAD2) lacked an SBS4 signature, showing profiles similar to non-smokers (Fig. 5B). One such tumour (LUAD1) showed prominent SBS5, SBS2 and SBS13 signatures (Fig. 5C). SBS5 is a ‘clock-like’ signature with some association with tobacco smoking, possibly due to the pathways involved rather than DNA damage by tobacco carcinogens. SBS5 shows a larger contribution to the former smoker samples than the non-smokers, although is present in both. These atypical mutational profiles are more similar to cancer types such as breast, rather than lung cancers (Fig. 5D). Note that to perform an analysis similar to the original Leong et al study (27) required extensive custom code.

#### Outcomes

Mutational signatures in this lung cancer cohort dataset confirmed most of these patients had tumours displaying profiles consistent with smoking-related mutagenesis. Two former-smoker patients, however, did not, and showed profiles suggestive of spontaneous oncogenesis. This work illustrates how CRUX can be used to streamline the analysis of mutation signatures, revealing unexpected findings.

### Short study 5 (cohort comparison mode): Gene mutations associated with triple-negative breast cancer

#### Rationale

An important subtype of invasive breast cancer with poor prognosis is triple negative breast cancer (TNBC), which lacks expression of *ER, PR*, and epidermal growth factor receptor 2 (*HER2*) genes. High levels of TP53 mutations have been reported in TNBC (28), making this a potential genetic biomarker. We investigated genes with mutations associated with TNBC status in breast cancer data using the CRUX ‘cohort comparison’ mode.

#### Dataset

We used the inbuilt TCGA Breast Invasive Carcinoma cohort dataset (n = 978) including ductal and lobular carcinomas, and filtered this dataset to n=969 females; the patient ages at first diagnosis were 26 to 90.

#### Analysis and observations

Clinical data provided by TCGA were used to construct a ‘TNBC’ cohort (n=104) of patient samples lacking *ER, PR* and *HER2* expression (i.e., ER^-^ and PR^-^ and HER2^-^), and a ‘not_TNBC’ cohort (n=757) of samples positive for at least one of these receptors (i.e., ER^+^ or PR^+^ or HER2^+^). To do this, the CRUX cohort subset feature was used on the following clinical data filters: ‘breast_carcinoma_estrogen_receptor_status’, ‘breast_carcinoma_progesterone_receptor_status’ and, ‘lab_proc_her2_neu_immunohistochemistry_receptor_status’ fields for ER, PR and HER2 status, respectively. For these filters only samples with values of ‘negative’ or ‘positive’ were used; those with values of ‘NA’ or ‘indeterminate’ were excluded. CRUX filtered out genes mutated in fewer than 5 tumour samples prior to cohort comparison. Forest plots, CoBar and CoOncoplots were used to compare mutation incidence between the groups, and lollipop plots to examine mutation location in genes of interest (Fig. 6).

**Figure 6.**
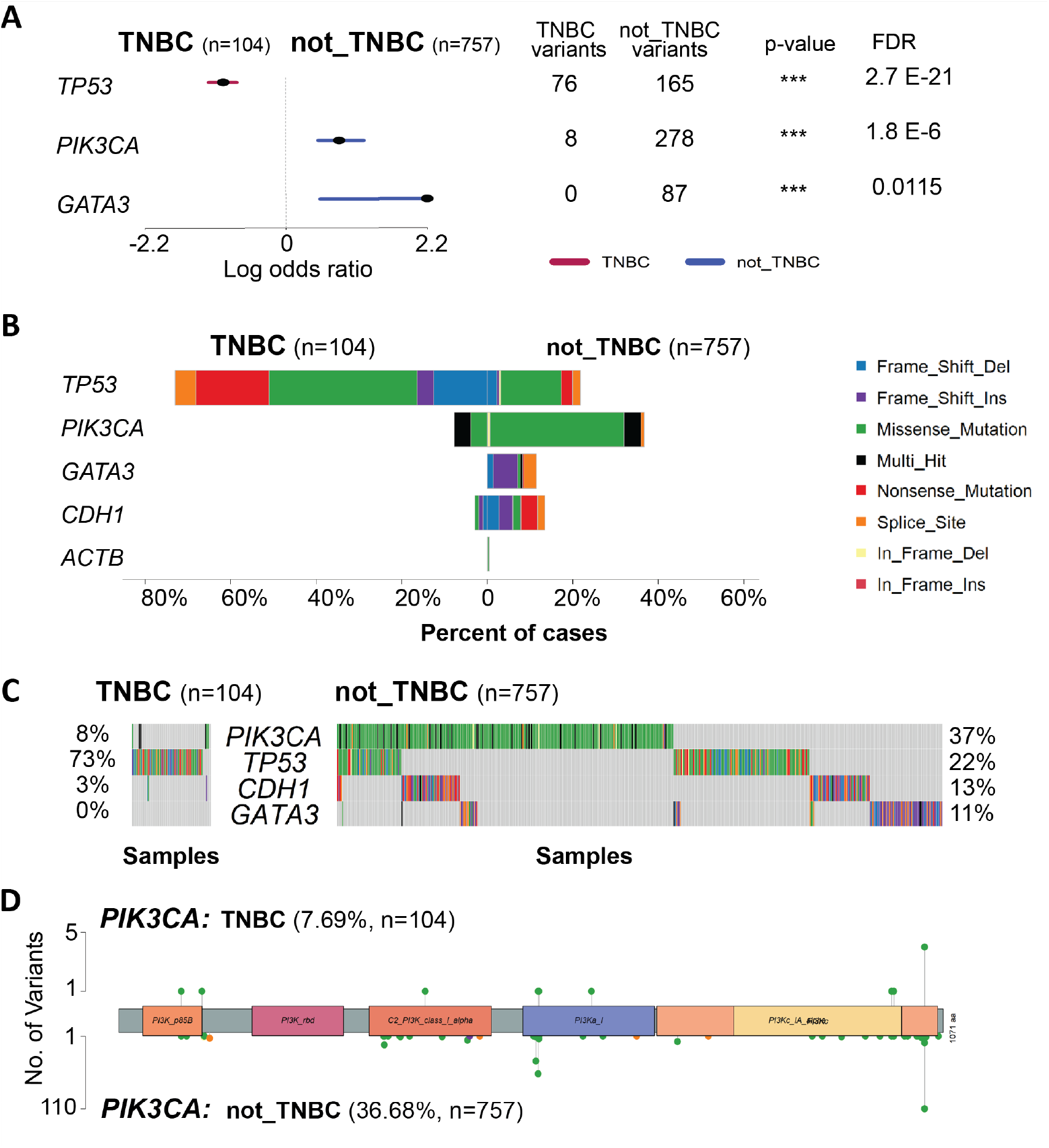
Differences in genetic variant incidence between TNBC and not_TNBC tumours from the TCGA Breast Invasive Carcinoma cohort. (A) Forest plot showing log odds ratio of variant incidence in *TP53, PIK3CA* and *GATA3* genes, which showed significant false discovery rate (<0.05) in the TNBC/not_TNBC comparison. Number of samples with variant incidence and unadjusted p-values for each gene are indicated, ***= p<0.001. (B) CoBar plot of most significant genes (plus ACTB, as negative control) showed differences in variant incidence between TNBC (left) and not_TNBC (right), with higher occurrence of *TP53* mutations in TNBC samples, and higher incidence of *PIK3CA, GATA3* and *CDH1* variants in non-TNBC tumours; note *CDH1* filtered out in A, as the as its relative incidence did not show FDR<0.05. Variant type is indicated, key on right. (C) CoOncoplots for gene variants in TNBC (left) and non-TNBC (right), where each column represents one tumour; colour key as in B. (D) Lollipop plot for *PIK3CA* gene, showing observed variant numbers for TNBC (top) and non-TNBC tumours (bottom); colour key as in B.

Forest plots comparing the mutation incidence between TNBC and not_TNBC (filtered using a false discovery rate of 0.05) identified three differentially mutated genes: *TP53, PIK3CA* and *GATA3* (Fig 6A). Consistent with these observations, CoBar plots (Fig. 6B) showed that missense and nonsense *TP53* mutations were enriched in TNBC samples vs not-TNBC (73% vs 23%). Conversely, *PIK3CA* and *GATA3* mutations showed lower incidence in TNBC (8% vs 37% and 0% vs 11%, respectively). Incidence of mutations in *CDH1* also appeared lower in TNBC (3% vs 13%), however this difference was not statistically significant (FDR > 0.05).

These genes have known associations with breast cancer. *PIK3CA* mutations are commonly found in these cancers and can predict responsiveness to PI3K inhibitors (29). *GATA3* transcription factor is expressed in breast ductal cells and crucial for breast development but, while there are many reports of *GATA3* mutations in breast cancer (including triple negative tumours), their functional effects are unclear (30). *CDH1*, a likely tumour suppressor gene, encodes cell adhesion molecule E-cadherin, with *CDH1* mutations (and abnormal methylation patterns) associated with invasive lobular breast carcinoma (31). The housekeeping gene, beta-actin (*ACTB)* was included as a negative control, revealing only rare mutations in either group (Fig. 6B). CoOncoplots (Fig. 6C) showed mutation distributions consistent with the CoBar plots, namely that most TNBC patients carry a TP53 mutation, and that non-TNBC patients carry combinations of one or more of these driver mutations. A lollipop plot of *PIK3CA* (Fig. 6D) showed that mutations were distributed across several protein domains, including the p.H1047R hotspot mutation in the N-terminus, with no apparent differences between the two cohorts. The house keeping gene, beta-actin (*ACTB)* was included as a negative control, revealing only rare mutations in either group (Fig. 6B).

*Outcomes*: Differences in mutation frequency of *TP53, PIK3CA and GATA3* between TNBC and not_TNBC breast cancers in women were evident using CRUX. This demonstrates how CRUX can be used to study possible novel molecular markers of cancer subtypes.

### Additional short study

An additional short study employing the CRUX single cohort mode to investigate copy number variants (CNV) in GBM and breast cancers from the pre-loaded TCGA datasets using the GISTIC2 tool is included in the Supplementary Results Section.

## DISCUSSION

The genomic analysis of cancer cohorts is revolutionising our understanding of cancer biology, enabling the identification of mutations that drive cancer development, providing insights into tumour formation, development, disease subtypes and treatment targets. Cohort analyses reveal not only cancer associated mutations and their frequencies, but also mutational signatures, gene amplifications, and fusion proteins. While a powerful research approach, a longstanding challenge is that exploring such tumour WGS data requires specialised analysis methods and algorithms, often beyond the capabilities of fundamental researchers. These methods are, however, increasingly available in high quality analytical packages that are often freely accessible, with many new packages developed each year. What is needed is a software tool that bridges the gap between users and those analysis packages.

This consideration led to our development of CRUX as a graphical environment for exploring mutations in cancer genomic data. We developed this platform using an R Shiny framework to give access to a range of functions and analysis tools through a user-friendly interface. This work builds on previous efforts to streamline cohort-level data exploration by extensively employing analysis tools and visualisations from the *maftools* R package (8). Since using *maftools* requires a deep knowledge of R, which greatly limits its user base, we developed CRUX to be a more accessible resource. CRUX thus facilitates easy use of *maftools* while also providing (in the same graphical environment) a consistent way to import publicly available and user-generated datasets, options to convert file formats, create and manipulate virtual cohorts, and run many types of analyses locally to mitigate data privacy concerns; CRUX also provides interoperability with many important web-based apps and platforms. Thus, CRUX makes many more analysis methods easily accessible and greatly enriches data analysis, making cohort-level exploration more time efficient. A previous approach to simplifying data access and analysis is TCGAbiolinksGUI (32). This resource simplifies TCGA data access with a graphical interface and methods to import genomic data commons resources (33), but has only limited genomic analytics capabilities. We have created the PCAWGmutations (9) and TCGAgistic R packages (34), which simplify the analysis of PCAWG mutation and TCGA CNV data, through precompiling these datasets, similar to the popular TCGAmutations (10) package; CRUX uses these packages to streamline access to this data.

Web-based platforms including cBioPortal (14), the St. Jude Cloud (1), and CVCDAP (5) also represent powerful analytical suites for cohort-level data. These platforms vary greatly in analysis methods that they provide, their diverse input file formats, support for analysis of public cancer datasets and ease of analysis of unpublished data (Table 1). However, with CRUX it is possible to directly leverage these resources, and many other specialised platforms. This is a particularly important design feature in CRUX given that no one package can offer all functionality, and that new platforms are constantly being developed and improved. This flexible approach means CRUX can keep analysis up to date with new advances in cancer genome analysis and visualisation tools.

To investigate the utility of CRUX to explore cancer mutation datasets we undertook several short studies which demonstrated that CRUX can be used to readily identify data features previously described in large-scale studies. For example, CRUX was able to employ a range of algorithms to identify *PTEN, EGFR* and *TP53* as the most likely drivers in a GBM cohort (35), while filtering many false positive hits (Fig. 2). CRUX can run identical analyses on any previously constructed or user-defined cohort to identify putative driver genes for any cohort of tumours. Similarly, lung cancers in smokers revealed a range of mutation profiles that give information on mutagenic exposures (Fig. 5). In another short study, mutations associated with increased survival (22) were identified in a GBM dataset (Fig. 3), and molecularly guided treatment options and drug sensitivities uncovered in a thyroid cancer cohort (Fig. 4). The genetic differences between phenotypically distinct breast cancer subtypes were also characterised (Fig. 6). While these short studies were not intended to break new ground or generate new insights, they all began with curiosity driven explorations and exploited the ability of CRUX to move quickly around a range of accessible modules. All these case studies highlighted discoveries that originally required extensive programmatic interrogation by genome bioinformaticians, capabilities that CRUX simplifies for all researchers and clinicians.

As part of such data explorations, users can construct virtual cohorts from imported datasets or from 57 pre-compiled public datasets (11,043 tumour samples), then analyse the cohort using any of the 19 modules available. A limitation of CRUX is that it lacks analysis modules to examine mutations in non-coding regions of the genome, nor ways to interrogate structural variants. As noted earlier, CRUX does not run the CNV analysis tool, GISTIC2 on custom user cohorts, but through our creation of the TCGAgistic R package (34), CRUX facilitates the seamless analysis of CNV data from TCGA. CRUX does not currently support the analysis of RNA-seq data, which is a future opportunity, given that these data are available for most of the tumours analysed by TCGA and PCAWG. Finally, we recommend that CRUX be used on a computer less than 5 years old with at least 8GB of RAM. For those users that would rather use a web-based version of CRUX, and are comfortable uploading their data to the internet, then we have made an online version of CRUX, available at https://ccicb.shinyapps.io/crux.

In summary, CRUX builds on previous efforts to leverage the use of powerful computational packages to enable cancer mutation exploration to be carried out by researchers and clinicians without a high level of training, which will help drive discovery research from cohorts of cancer patients.

## DATA AVAILABILITY

TCGA datasets packaged with CRUX, including the GBM (36), THCA (37) and BRCA (38) datasets analysed in the short studies presented here, were sourced from the TCGAmutations package (10). PCAWG datasets (7) were sourced from the PCAWGmutations package (9) available at https://github.com/CCICB/PCAWGmutations, DOI: 10.5281/zenodo.8115522. Lung carcinoma datasets used for mutational signature analysis were published in Leong et al. (2019) available from the European Genome-Phenome Archive under accession number EGAD00001005287 (27). VCFs were converted to MAFs using vcf2maf. Broad Institute GISTIC2 data for BRCA (39) and GBM (39) cohorts were compiled from Firebrowse, 2016-01-28 analysis (39), and streamed into R using the TCGAgistic package (34) available from https://github.com/CCICB/TCGAgistic, DOI: 10.5281/zenodo.8115633

## SUPPLEMENTARY DATA

Supplementary data is available in ‘supplementary_data.pdf’

## ACKNOWLEDGEMENTS

The results published here are in whole or part based upon data generated by the TCGA Research Network (https://www.cancer.gov/tcga) and the PCAWG resource generated by the International Cancer Genome Consortium (https://docs.icgc.org/pcawg), as described above.

