## Supplementary Materials for "CRUX, a platform for visualising, exploring and analysing cancer genome cohort data"

### SUPPLEMENTARY RESULTS

#### <u>Additional Short Study (single cohort mode)</u>: Identifying key copy number variants (CNVs) in a tumour cohort dataset

*Rationale*: Somatic CNVs are a hallmark of many tumours, driving large-scale numerical differences in DNA, resulting in alterations to gene dosage and expression. GISTIC2 is a well-established tool for assessing the CNV landscape of a cancer cohort, identifying genomic regions harbouring recurrently amplified and deleted genes. CRUX can use GISTIC2 processed data (which are publicly available for TCGA datasets) to visualise CNV and add clinical detail annotations for further interactive explorations. To keep CRUX lightweight and easy to install, CRUX does not run GISTIC2 itself, and instead offers pre-compiled GISTIC2 data for TCGA cohorts. For researchers wishing to generate GISTIC2 results on their cohort, we recommend using GenePattern (available online at www.genepattern.org).

*Dataset*: Inbuilt TCGA GISTIC2 results for both GBM (n = 273 with matched WGS) and breast carcinoma (n=960 with matched WGS) were used.

*Analysis and Observations*: Here, the TCGA GISTIC2 data was imported into CRUX, then the GISTIC-Plot module was used to create tabulated summaries and data visualisations highlighting potentially important CNVs (Supplementary Fig. 5A). The chromplot visualisation (Supplementary Fig. 5B) revealed the highest G-Score peak, a measure of CNV amplitude and recurrence, which corresponded to *EGFR* amplification in GBM; the second strongest CNV signal was a deletion in the *CDKN2A*-containing 9p21 region. Four other smaller peaks were noted with G-scores between +0.5 and +2. An analysis of the TCGA breast carcinoma cohort CNV data showed strong signals of *HER2* / *ERBB2* amplification, well described in breast carcinoma (Supplementary Fig. 5C).

*Outcomes*: CRUX visualised several CNV events that are known drivers of cancer development, including CNVs affecting *EGFR* in GBM and *ERBB2* (HER2) in breast cancers.

### SUPPLEMENTARY TABLES

| Property | TCGA Biolinks GUI | cBio Portal | St. Jude Cloud | Gene Pattern | CVCDAP | Maf tools | CRUX |
|---|---|---|---|---|---|---|---|
| **Inbuilt support for public data analysis** | ✓ | ✓ | ✓ | ✓ | ✓ | ✓ | ✓ |
| **Supports analysis of unpublished user data** | ✓ | ✓ (orange) | ✓ (orange) | ✓ | ✓ | ✓ | ✓ |
| **Runs locally (Windows/ MAC)** | ✓ | | | | | ✓ | ✓ |
| **Dedicated inter-operability module** | | | | | | | ✓ |

**Supplementary Table S1. Key features of cancer cohort-level analysis suites.** Green tick symbol: key features supported. Orange tick symbol: functionality is technically supported but with at least one of the following caveats: 1) technically demanding creation of a local instance, 2) resource maintainers need to be contacted to organise integration of user data, 3) upload of data to paid cloud services, or 4) code needs to be written or run by end-users.

| Class | Type of Variation | Analysis / Visualisation |
| --- | --- | --- |
| Single Cohort Analyses | SNVs and Indels | Oncoplot |
| | SNVs and Indels | TiTv Rates |
| | SNVs and Indels | Highly Mutated Pathways |
| | SNVs and Indels | Gene level lollipop plots |
| | SNVs and Indels | Pfam Domain Mutation rates |
| | SNVs and Indels | Somatic Interactions (gene-level cooccurrence of mutations) |
| | SNVs and Indels | Survival Analysis for prognostic biomarker identification |
| Enrichment Analysis | SNVs and Indels | Genomic Enrichment Analysis |
| Two-Cohort Analysis | SNVs and Indels | Cohort Comparison |
| Single Cohort | CNVs | GISTIC2 Chromplot |
| | | GISTIC2 Oncoplot |
| Sample Level | SNVs | Kataegis |
| Sample Level | Heterogeneity | Heterogeneity estimation |
| Utilities | SNVs and Indels | Subset / Merge Cohorts |

**Supplementary Table S2: Modules available in CRUX**

#### SUPPLEMENTARY FIGURES

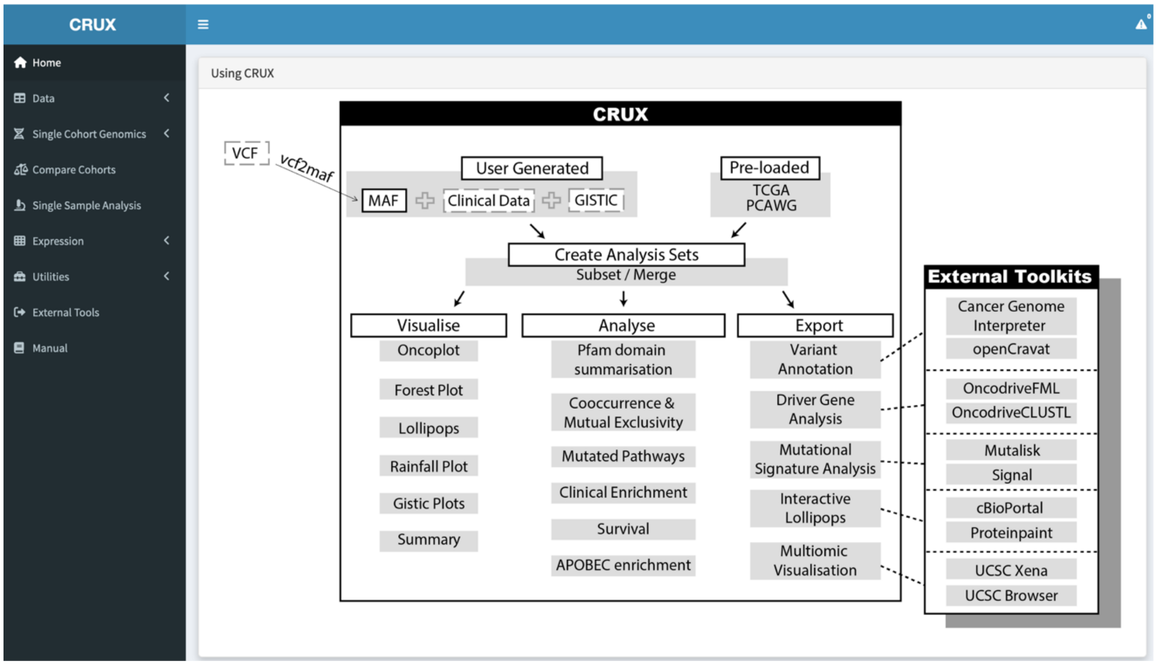


**Supplementary Figure 1. CRUX landing page**. This screenshot of the CRUX user interface window shows the available workflows, the organisation of tools into visualisation, analysis and export tools for using several external toolkits.

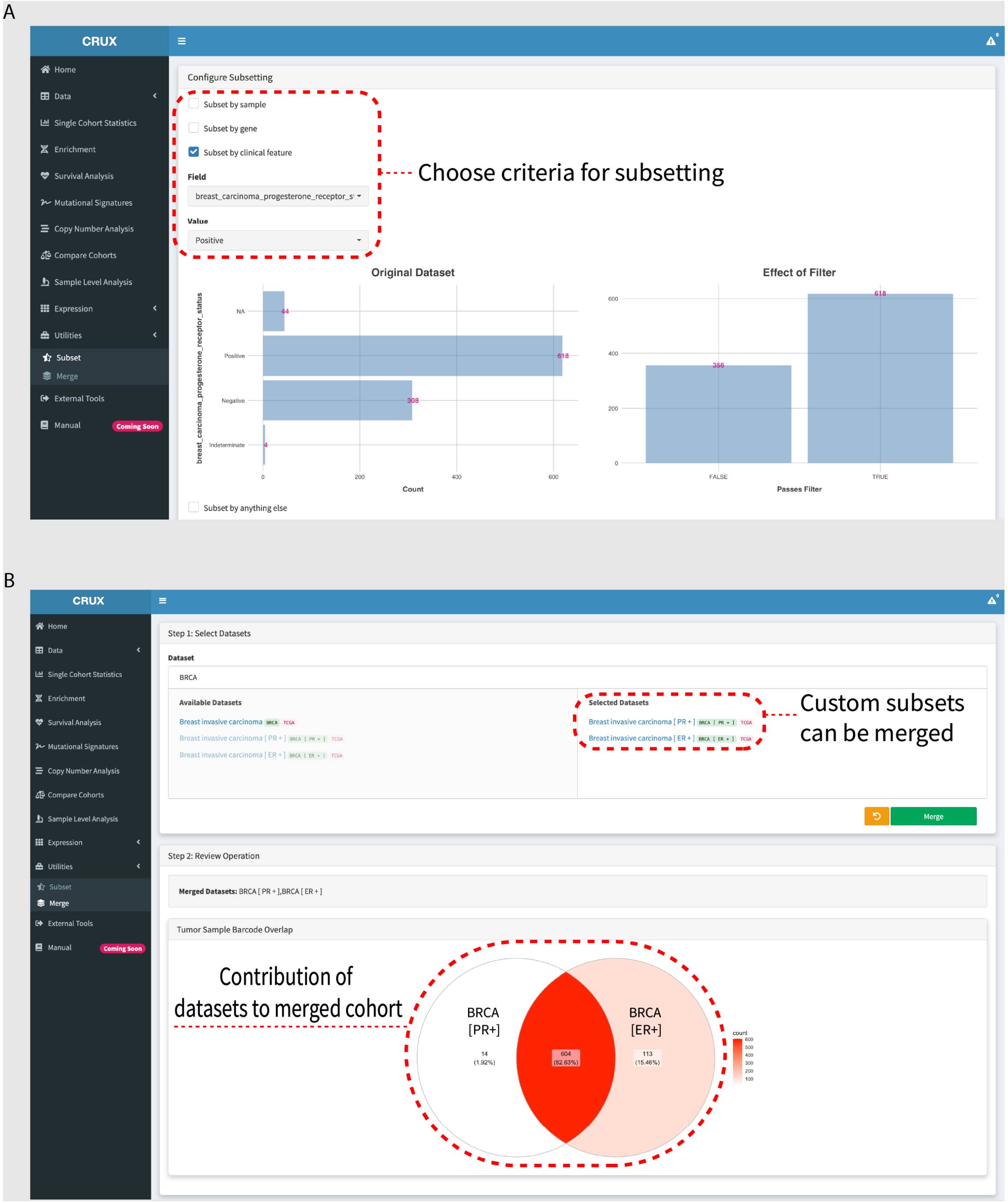


**Supplementary Figure 2**. **Creation of new cohorts in CRUX**. This is illustrated by screenshots of CRUX user interface window. (A) Subsetting and (B) merging of cohorts to create new analysis sets. Subsetting can be based on molecular abberations or sample-level metadata.

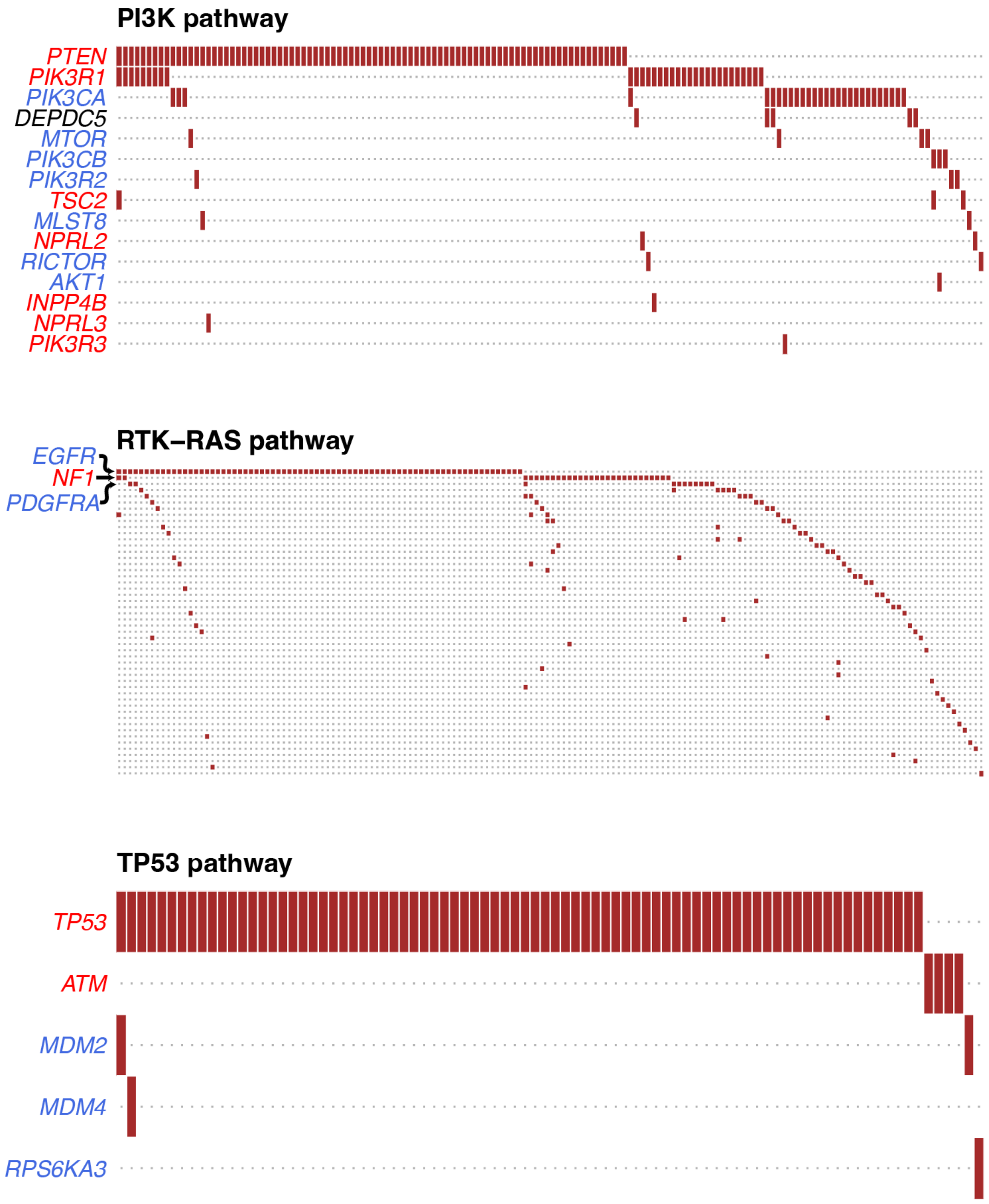


**Supplementary Figure 3**. Gene-level pathway analysis for highly mutated pathways. CRUX facilitates more granular investigation of individual pathways. Columns represent individual samples, with each coloured box representing a that the given gene is mutated in the sample. Gene names are coloured red for tumour suppressor genes and blue for oncogenes.

| Sample Id | Gene | Protein change | Consequence | Domain | *Oncogenic classification* | GDNA |
|---|---|---|---|---|---|---|
| Search here... | TTN | | | | | |
| TCGA-DJ-A13O | TTN | p.L21223F | Missense | fn3 | predicted passenger | chr2:g.179439488G>A |
| TCGA-DJ-A1QO | TTN | p.E7343D | Missense | I-set | predicted passenger | chr2:g.179580380C>G |
| TCGA-DJ-A2Q3 | TTN | p.R27041* | Nonsense | fn3 | predicted passenger | chr2:g.179419249G>A |
| TCGA-DJ-A3VA | TTN | p.P441Q | Missense | Titin_Z | predicted passenger | chr2:g.179659202G>T |
| TCGA-E3-A3E1 | TTN | p.T18177Rfs*10 | Frameshift | fn3 | predicted passenger | chr2:g.179454219_17… |
| TCGA-E8-A417 | TTN | p.V13170L | Missense | fn3 | predicted passenger | chr2:g.179482973C>G |
| TCGA-EL-A3CU | TTN | p.T22056I | Missense | fn3 | predicted passenger | chr2:g.179436988G>A |
| TCGA-EL-A3MY | TTN | p.Y30639C | Missense | | predicted passenger | chr2:g.179402314T>C |
| TCGA-EM-A3AR | TTN | p.I28777M | Missense | fn3 | predicted passenger | chr2:g.179412318T>C |
| TCGA-ET-A25K | TTN | p.K17826Q | Missense | fn3 | predicted passenger | chr2:g.179455272T>G |
| TCGA-IM-A3U3 | TTN | p.P3924T | Missense | I-set | predicted passenger | chr2:g.179598614G>T |
| TCGA-EL-A4K6 | TTN | p.R14198* | Nonsense | fn3 | predicted passenger | chr2:g.179476842G>A |
| TCGA-EM-A3ST | TTN | p.E20463* | Nonsense | I-set | predicted passenger | chr2:g.179441971C>A |
| TCGA-EL-A4KI | TTN | p.R3257C | Missense | I-set | predicted passenger | chr2:g.179629473G>A |
| TCGA-EL-A3ZS | TTN | p.S19050F | Missense | fn3 | predicted passenger | chr2:g.179449515G>A |
| TCGA-EL-A3ZS | TTN | p.P24898H | Missense | | predicted passenger | chr2:g.179428462G>T |
| TCGA-EL-A3TB | TTN | p.M31494T | Missense | Pkinase | predicted passenger | chr2:g.179399157A>G |
| TCGA-E8-A432 | TTN | p.W31431R | Missense | Pkinase | predicted passenger | chr2:g.179399347A>T |
| TCGA-BJ-A0ZB | TTN | p.Y29510Y | Synonymous | I-set | not protein-affecting | chr2:g.179408637G>A |
| TCGA-DE-A4MC | TTN | . | IntronicSNV | | not protein-affecting | chr2:g.179610827C>T |
| TCGA-EL-A3H7 | TTN | p.S7441S | Synonymous | I-set | not protein-affecting | chr2:g.179579858G>A |
| TCGA-EM-A4FN | TTN | . | IntronicSNV | | not protein-affecting | chr2:g.179611913T>G |
| TCGA-ET-A3BW | TTN | . | IntronicSNV | | not protein-affecting | chr2:g.179610555A>G |
| TCGA-EL-A3CS | TTN | . | IntronicSNV | | not protein-affecting | chr2:g.179612467A>G |

**Supplementary Figure 4**. Cancer Genome Interpreter (CGI) annotation of Titin (*TTN*) variants present in the TCGA thyroid cancer cohort. The most severe oncogenic classification of *TTN* variants is 'predicted passenger', which indicates *TTN* mutations are very unlikely to drive thyroid cancer development, so *TTN* cannot be classified as a tumour driver gene.

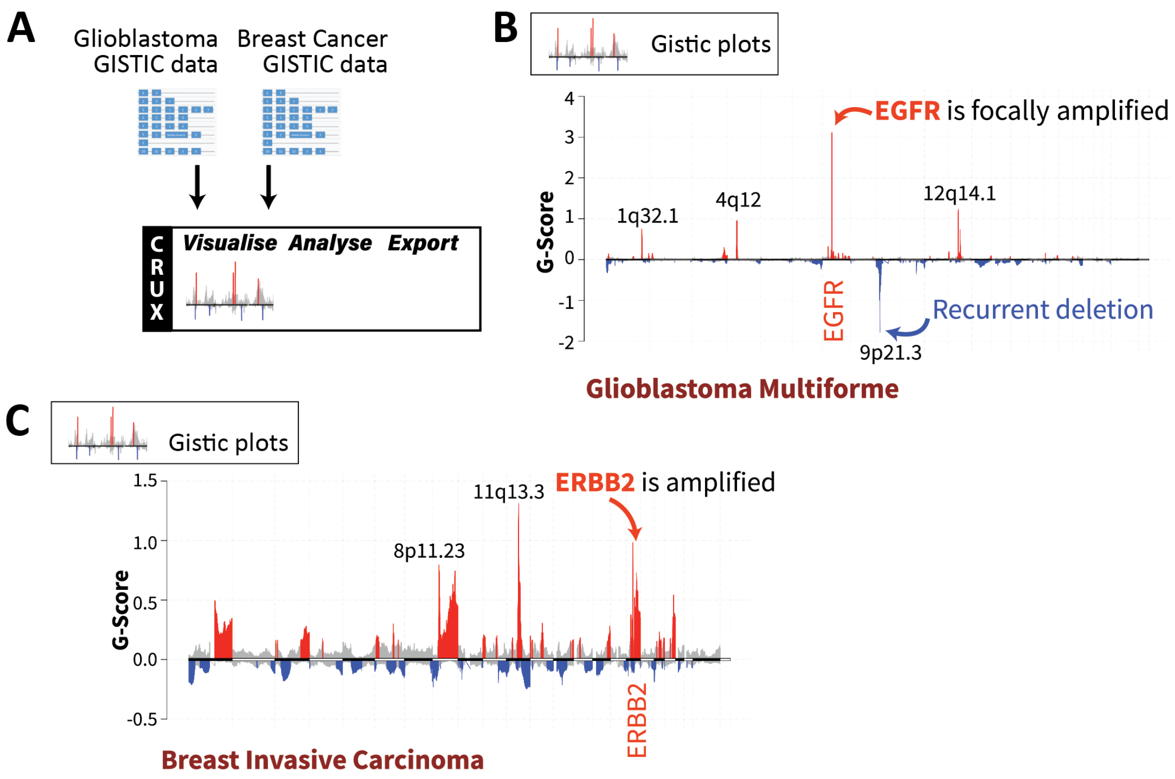


**Supplementary Figure 5. Cohort-level visualisation of copy number alterations from GISTIC2 results using CRUX.** (A) Integration of CNV data from TCGA glioblastoma (GBM) and TCGA breast carcinoma (BRCA). Inbuilt GISTIC2 datasets for the TCGA GBM and BRCA cohorts were selected, leading to automatic generation of 'chromplot' visualisations. These are plots of the G-Scores (a metric integrating CNV recurrence and amplitude) which indicate the likely importance of CNVs in a cohort. (B) Selecting the GBM cohort in CRUX produced visualisations showing peaks corresponding to the characteristic EGFR amplification. (C) Visualisation of invasive breast carcinoma CNV data reveals peaks that notably correspond to HER2 / ERBB2 as well as 11q13.3 amplification, which contains driver genes such as CCND1 and EMS1.
